# Pan-cancer analysis reveals the value of FDX1 as a novel biomarker in the prognosis and immunotherapy in human tumors

**DOI:** 10.1101/2022.06.22.497145

**Authors:** Qingqing Zhang, Nanyang Liu, Di Wu, Zhengyu Xu, Yichen Wang, Ping Wang

## Abstract

Emerging evidence supports FDX1’s important role in the development and progression of cancer, but no extensive cancer analysis is available to date. This study was the first to comprehensively explore the expression of FDX1 in 33 types of cancer and the significance of FDX1 in clinical prognosis by using bioinformatics techniques. Meanwhile, we analyzed the relationship between FDX1 and pathological stage, as well as immune cell infiltration. Based on this, the important role of FDX1 in tumor immunotherapy was proposed. The expression of FDX1 was significantly different between normal and tumor samples in 17 of 33 types of cancer. Besides, Cox regression analysis showed that FDX1 is a protective gene in Kidney renal clear cell carcinoma (KIRC), Cervical squamous cell carcinoma and endocervical adenocarcinoma (CESC), Liver hepatocellular carcinoma (LIHC), Kidney renal papillary cell carcinoma (KIRP), Mesothelioma (MESO), and Thyroid carcinoma (THCA). Porphyrin and oxidative metabolism pathway regulating integrator complex was involved in the process. Furthermore, high expression of FDX1 promoted infiltration of Eosinophils and monocyte in Adrenocortical carcinoma (ACC) and Kidney Chromophobe (KICH) by affecting the tumor microenvironment (TME) and was significantly correlated with immune checkpoint genes. Our first pan-cancer analysis elucidates the expression characteristics of FDX1 across different cancers and highlights its potential value as a prognostic biomarker, laying a foundation for further study of its immunotherapy mechanism in various cancers.

## Introduction

In the wake of increasing amounts of new cancer cases every year, cancer immunotherapy (CIT) focused on T cells plays an essential role in the armamentarium against cancer^[1]^. Although the treatment has been widely used in clinical practice, most cancer patients respond poorly to this treatment^[2]^. Thus, it is urgent to explore a new biomarker with predictive function to assess the response to immunotherapy from patients, which helps us identify benefits to patients early.

Metabolism takes a vital part in carcinogenesis, and multiple studies have clarified that metabolism and cancer are closely related ^[3, 4]^. As the research progresses, it has gradually been leveraged for cancer treatments, which metabolites are regarded as indispensable targets in the treatment of cancer^[5]^. Nutrient and oxygen deficiency, ATP depletion, and inflammatory response in the tumor microenvironment (TME) create a long-term stressful environment for tumor cells to live in. It is known that immune cells are one of the tissue cells highly dependent on oxidative metabolism. Fatty acid metabolism of immune cells is a critical metabolic process in the tumor microenvironment. For example, fatty acid synthesis(FAS) provides raw materials for the proliferation of effector T cells and facilitates functional maturation of Treg cells, while fatty acid oxidation (FAO) inhibits the activation of effector T cells and reduces the secretion of interferon-γ^[6]^. Increased FAS in tumor cells and gradual accumulation of fatty acids in the tumor microenvironment induce effector T cell senescence, promoting tumor progression.

Strengthening FAO can promote the differentisation of memory T cells in the TME and is favorable for antigen reactivation. Meanwhile, corticosteroids may interfere with the fatty acid metabolism and reduce the number of low-affinity memory T cells, inhibit the proliferation of T cells, and weaken the anti-tumor effect of cytotoxic T lymphocyte-associated antigen 4 monoclonal antibody^[7]^.

FDX1, a member of the [2Fe-2S] cluster-containing ferredoxin family, encodes a small iron-sulfur protein involved in the reduction of mitochondrial cytochromes and the synthesis of various steroid hormones^[8]^. More and more evidence suggests that FDX1 can promote the synthesis of various steroid hormones, and ATP production, and the knockdown of FDX1 significantly affected glucose metabolism, fatty acid oxidation, and amino acid metabolism^[9]^. Furthermore, the fact that the high expression of FDX1 contributes to the death of tumor cells by affecting the process of metabolism and suppressing the development of cancer has been confirmed in recent research^[10-13]^. Therefore, regulating the oxidative metabolism of the tumor microenvironment to inhibit the growth and promote the death of tumor cells by commanding the expressive level of FDX1 is of great significance for controlling the progress of cancer and strengthening the effect of immunotherapy.

Whereas studies on the potential of FDX1 as a predictive biomarker of immunotherapy response in various cancer types remain unexplored. In this paper, we performed a pan-cancer analysis of the FDX1 gene across 33 cancer types using the Cancer Genome Atlas (TCGA) dataset. The expression of FDX1 was significantly different between normal and tumor samples in 17 of 33 types of cancer. Subsequently, the association of FDX1 with patient survival and tumor progression in these 17 cancers was investigated. Low expression of FDX1 was significantly associated with poor prognosis in several cancers. The ability of FDX1 to predict tumor immunity was also explored. High expression of FDX1 promoted infiltration of Eosinophils and monocyte and was strongly correlated with the immune checkpoint. Our pan-cancer analysis elucidates the expression characteristics of FDX1 in different cancers and highlights its potential value as a prognostic biomarker, laying a foundation for further study of its immunotherapy mechanism in various cancers.

## Methods

### Data Processing and Differential Expression Analysis

RNA sequencing, somatic mutation, and related clinical data were downloaded from TCGA (which contains 11069 samples from 33 types of cancer) using UCSC Xena (https://xena.ucsc.edu/), an online tool for the exploration of gene expression, clinical, and phenotype data. Strawberry Perl (Version 5.32.0, http://strawberryperl.com/) was used to extract FDX1 gene expression data from these downloaded data sets and plot it into a data matrix for subsequent analyses. A comprehensive pan-cancer analysis was performed to clarify its aberrant expression across 33 cancer types using the downloaded data and expression levels of 33 tumor samples compared with matched standard samples. Log2 transformation of expression data was performed, and two t-tests were performed for these tumor types. P <0.05 was the difference of expression between tumor and normal tissue. The r package “ggpubr” was utilized to draw box plots using R version 4.2.0 software.

### Clinical correlation analysis

The survival data downloaded from TCGA were analyzed for the clinical correlation with FDX1 in a variety of cancer, and the expression of FDX1 in different ages, gender, and tumor stages was obtained. P< 0.05 was considered to have differential expression of FDX1 in different stages. Data analysis was conducted using R software (Version 4.0.2;), and the R package “ggpubr” was used to draw box plots.

### Survival analysis

Survival analysis files were obtained from FDX1 expression files and survival data downloaded from TCGA for 33 tumors using r software. The relationship between FDX1 expression and patient prognosis involving overall survival (OS), disease-free survival (DFS), disease-specific survival (DSS), and progression-free survival(PFS) was obtained in the following two ways. First, the samples were divided into high and low expression groups according to the median expression of FDX1, and a Kaplan-Meier analysis was performed to compare the survival difference between the two groups. The other is to compare the relationship between FDX1 expression and survival time and survival state by the Cox proportional-hazards model. Any differences in survival were evaluated with a stratified log-rank test. P < 0.05 as significant. In the Cox analysis, FDX1 was divided into high-risk (HR>1) and low-risk genes (HR<1) according to the relationship of HR value to 1, and then the HR value was visualized, and a forest plot was drawn using the “forest plot” package.

### Gene Set Enrichment Analysis (GSEA)

To explore the functions and pathways active in the high-level FDX1 group of each cancer providing the evidence with respect to possible biological functions of FDX1 in various cancers, we performed Gene Set Enrichment Analysis (GSEA) based on the downloaded gene sets c2.cp.kegg.v7.4.symbols.gmt and c5.go.v7.4.symbols.gmt using the R-packages “colorspace”, “string”, “ggplot2”(P < 0.05 as significant).

### Differential analysis of the gene activity in pan-cancer

FDX1 based on single-sample gene set enrichment analysi(s ssGSEA). Using the sorted expression files of 33 tumors and the merged pan-cancer expression files, the gene most related to FDX1 was found by the co-expression method. That is, the active genes of FDX1, and use ssGSEA^[14]^ to score the active genes. Finally, the distribution situation defines tumors as high and low expression groups (the upper part corresponds to the high group, and the lower part corresponds to the low group). The boxplots concerning the differential analysis of the gene activity were obtained using R the “ggpubr” package.

### Analysis of the relationship between FDX1 expression and immunity

Tumor microenvironment (TME) cells are important elements that make up tumor tissue and can assess the therapeutic effects. In this paper, the immune and stromal scores for each tumor sample were calculated using the ESTIMATE^[15]^ algorithm regarding the downloaded FDX1 expression data and the correlation based on the correlation coefficient of 0.3 between FDX1 expression and these two scores assessed by the degree of immune invasion. Moreover, for obtaining the content of immune cells in each sample, the “CIBERSORT” was used to calculate immune infiltration. In this process, the R software packages “ggpubr” and “ggplot2” was used to visualize the results (P < 0.05 as significant).

In addition, we further analyzed the co-expression of FDX1 and immune-related genes and downloaded relevant data on the website(http://cis.hku.hk/TISIDB/), including genes encoding major histocompatibility complex (MHC), immune activation, immunosuppressive.

### Correlation of FDX1 expression with Tumor Mutation Burden and Tumor Microsatellite Instability

Tumor Mutation Burden (TMB) is a quantifiable immune response biomarker that reflects the total number of somatic gene coding errors, base substitutions, gene insertion or deletion errors^[16]^, which results in Microsatellite Instability(MSI). The related scripts were run in Perl to obtain tumor mutation coincidence files that were analyzed to obtain correlation with FDX1 expression level. Microsatellite Instability (MSI) scores were determined for all samples based on somatic mutation data downloaded from TCGA (https://tcga.xenahubs.net) and the relationship between FDX1 expression and TMB and Microsatellite Instability MSI was analyzed. All the above data were analyzed using R-package “fmsb” and the results were shown by a radar chart.

### Statistical Analysis

The gene expression data were transformed by LOG2, and a t-test was used for comparison between normal tissues and cancer tissues. Kaplan-Meier was used to investigate the correlation between FDX1 expression level and patient survival time. Cox proportional risk regression model was used to calculate HR and P values for survival analysis, and the results were analyzed by Spearman or Pearson test. In all analyses, P < 0.05 was considered statistically significant and was processed by R software (version 4.2.0).

## Results

### Differences in FDX1 expression between tumor and normal tissues

Our study found that FDX1 was expressed in all tumors (Figure 1A), with the highest levels in Adrenocortical carcinoma (ACC) and the lowest in Uveal Melanoma (UVM). Subsequently, differences in FDX1 expression between tumor and normal samples were known (Figure 1B). Except for tumors that did not contain normal samples in TCGA, there was a significant difference concerning the expression of FDX1 between tumor tissues and normal tissues in 17 types of cancer, including Breast invasive carcinoma(BRCA), Cervical squamous cell carcinoma, and Cholangiocarcinoma(CHOL), Colon adenocarcinoma(COAD), Glioblastoma multiforme (GBM), Head and Neck squamous cell carcinoma(HNSC), Kidney Chromophobe (KICH), Kidney renal clear cell carcinoma(KIRC), Kidney renal papillary cell carcinoma(KIRP), Liver hepatocellular carcinoma(LIHC), Lung adenocarcinoma(LUAD), Lung squamous cell carcinoma (LUSC), Pheochromocytoma and Paraganglioma (PCPG), Rectum adenocarcinoma (READ), Sarcoma (SARC), Stomach adenocarcinoma (STAD), Thyroid carcinoma (THCA), Uterine Corpus Endometrial Carcinoma (UCEC). It is worth mentioning that compared with normal tissues, FDX1 expression was down-regulated except in Stomach adenocarcinoma (STAD) and Uterine Corpus Endometrial Carcinoma (UCEC), and the extent of FDX1 reduction is most dramatic in Pheochromocytoma and Paraganglioma (PCPG). In addition, the reason that the tumors with no significant difference in expression may be due to the small number of normal samples or the absence of corresponding normal samples in the TCGA database. Meanwhile, the analysis of FDX1 activity illuminated that FDX1 had the lowest activity in the Skin Cutaneous Melanoma (SKCM) (Figure 1C), highest activity in the Lymphoid Neoplasm Diffuse Large B-cell Lymphoma (DLBC), possess discrepancy outstandingly between normal and tumor tissues in 16 cancers, and decline noteworthily compared with the normal tissue (Figure 1D).

**Figure1.**
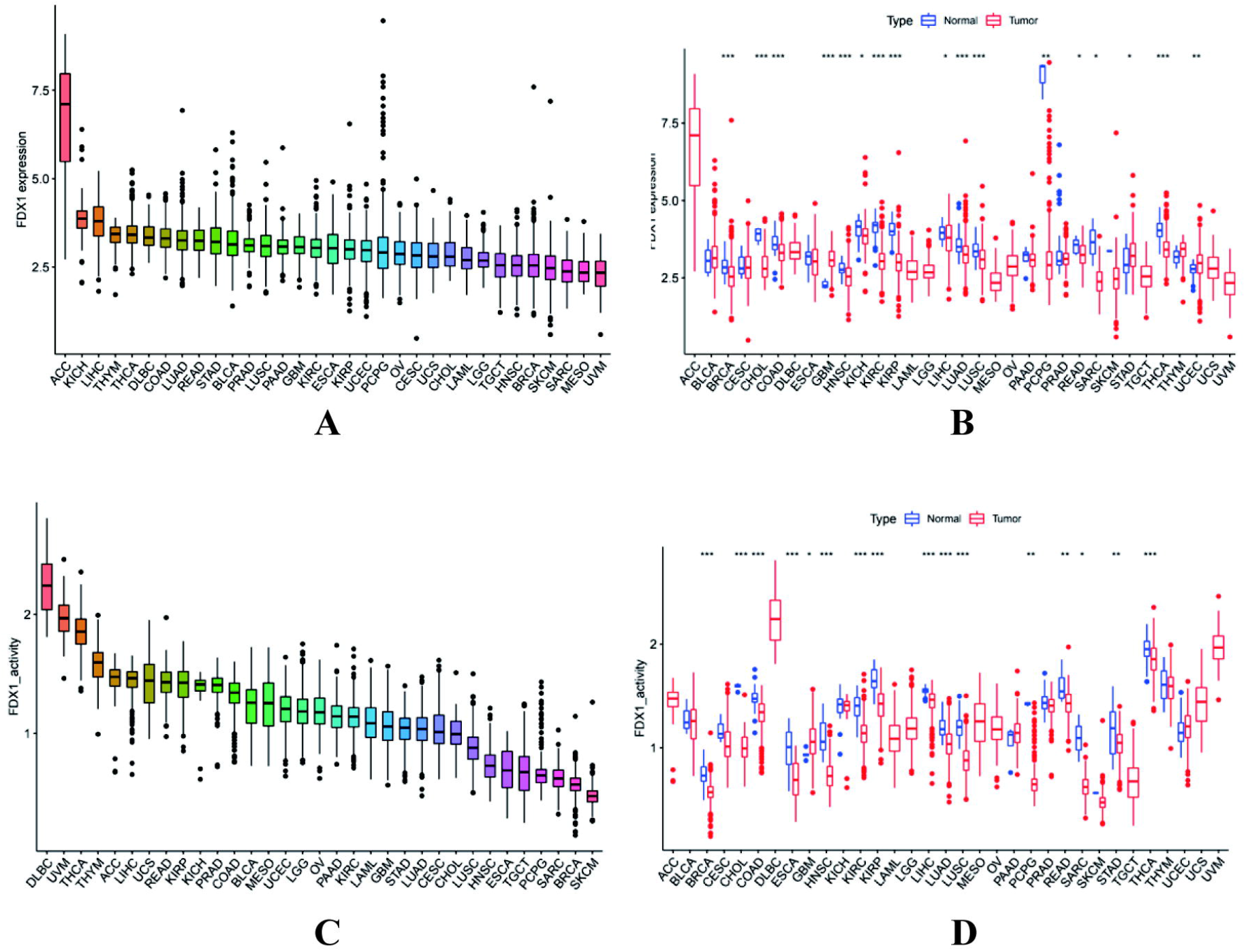
The expression of FDX1. **(A)** The expression of FDX1 in 33 tumors **(B)**The differences in FDX1 expression between tumor and normal tissues **(C)**The activity of FDX1 in 33 tumors **(D)**The differences in gene activity of FDX1 in 33 tumors. p<0.05 was considered significant, *p<0.05, **p<0.0l, ***p<0.001

### Correlation between FDX1 expression and clinical phenotype in pan-cancer

For the sake of exploring the relationship between FDX1 expression and clinical phenotype in 33 cancers, the correlation analysis with FDX1 on the data downloaded from the TCGA database involving the gender, age, and tumor stage of patients was performed. The results clarify that FDX1 expression had no significant correlation with age and sex in most cancers. Among age and gender-related tumors (P <0.5, ESCA, HNSC, UCEC and BRCA, KIRC), FDX1 expression was higher in patients older than 60 years of age and male patients (Figure2A-B). In contrast, the level of FDX1 was significantly associated with tumor stage in 4 cancers, including ESCA, KICH, LIHC, and THCA, and gradually down-regulated from stage 1 to stage 3 (Figure2C).

**Figure2.**
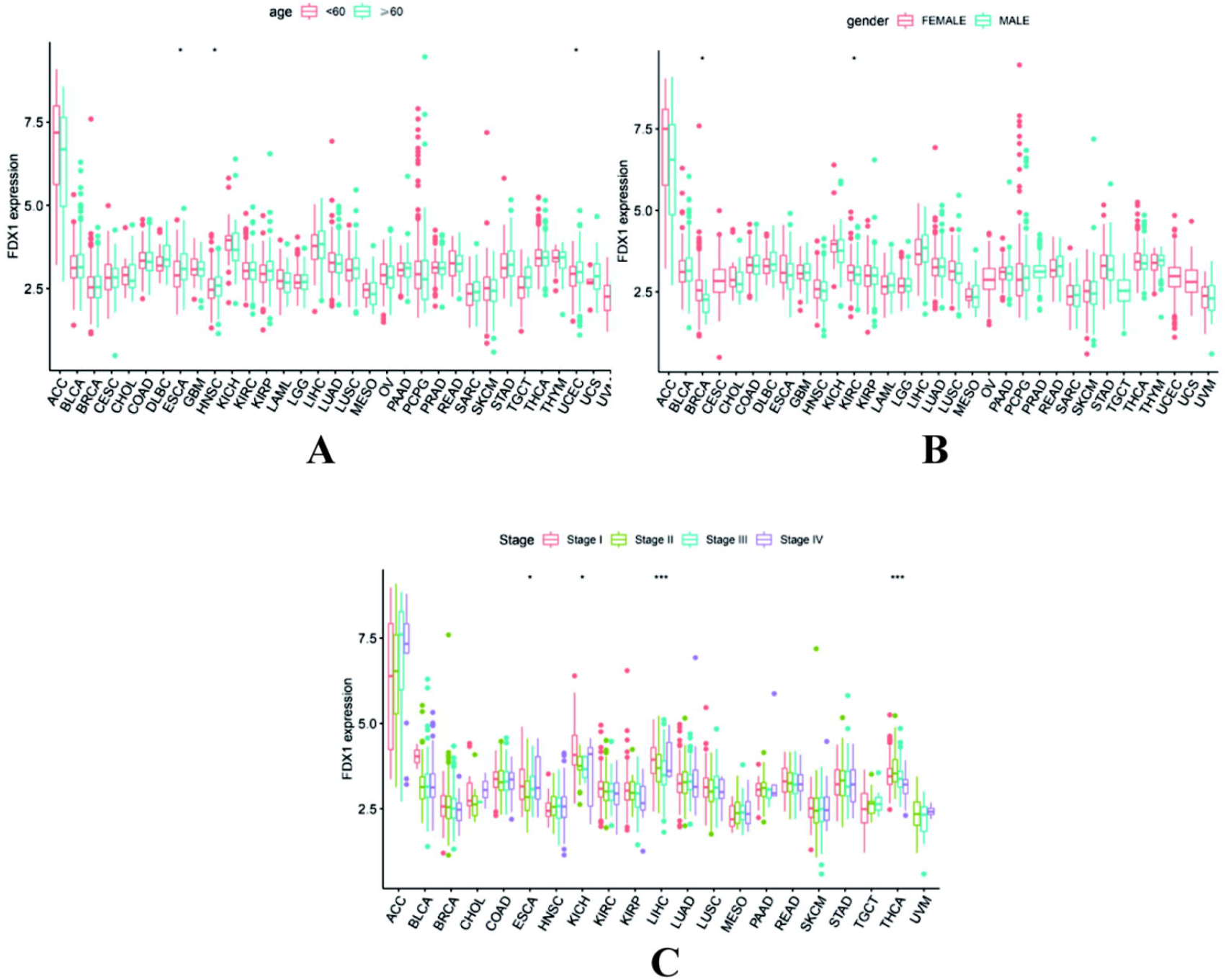
Correlation between FDX1 expression and clinical phenotype in 33 cancers **(A)**Differential expression of FDX 1 in ages **(B)**Differential expression of FDX 1 in genders **(C)**Differential expression of FDX 1 in tumor stages. p<0.05 was considered significant, *p<0.05, **p<0.0l, ***p<0.001

### FDX1 expression with patient prognosis and GSEA in pan-cancer

Cox regression analysis showed that FDX1 is a protective gene in KIRC, CESC, LIHC, KIRP, MESO, and THCA(Figure3). The Kaplan-Meier curves between high expression and low expression samples indicated FDX1 expression was associated with the patient’s prognosis in 11 types of cancer, including ACC, COAD, HNSC, KIRC, LGG, PAAD, SKCM, KICH, LIHC, THCA, MESO. Among these cancers, the low expression group had a poor prognosis in COAD(OS), KIRC (OS, DSS, PFS), SKCM(OS), THCA (DFS, PFS), LIHC (DFS, DSS, PFS), and MESO(DSS) (Figure4).

**Figure3.**
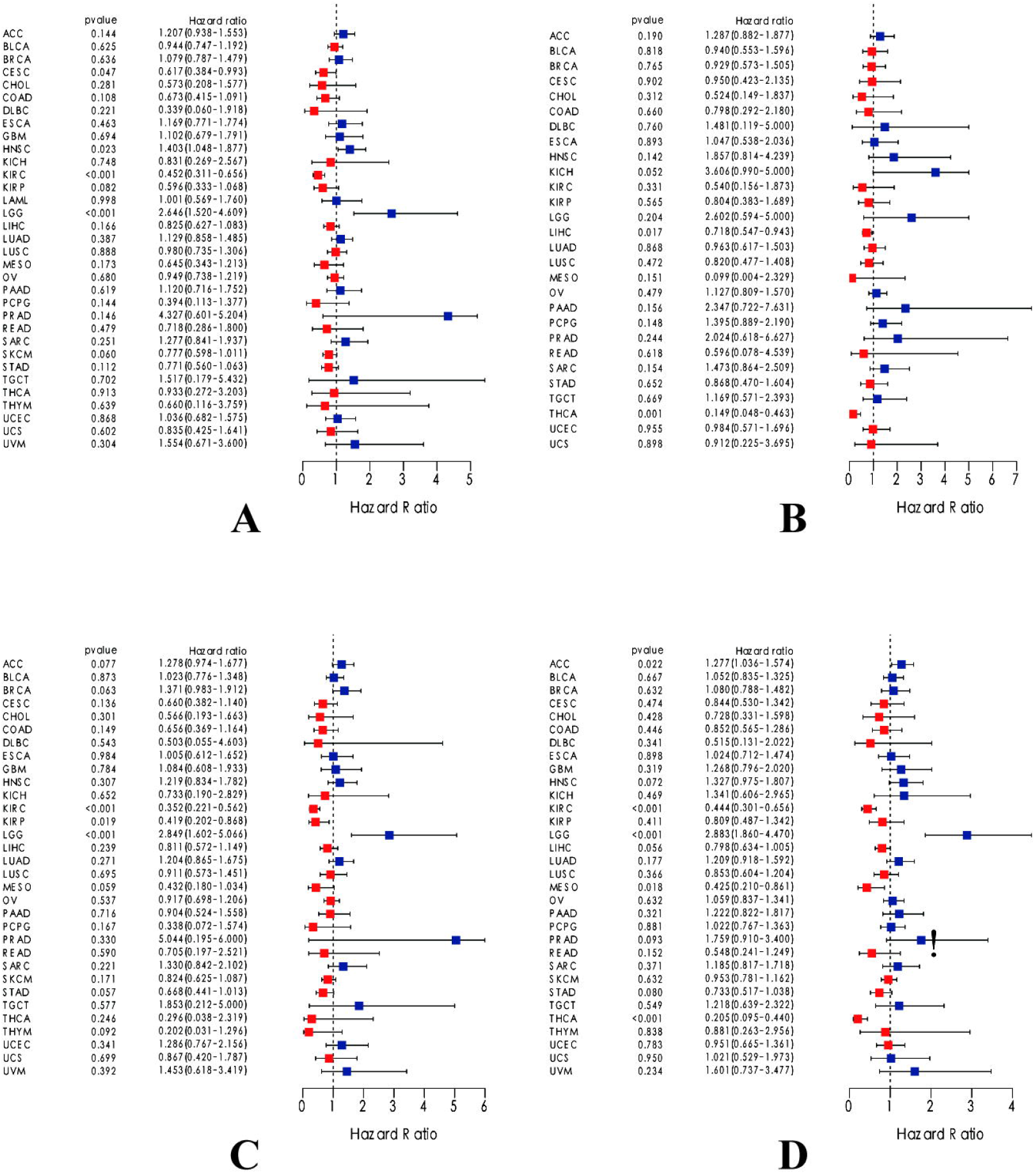
Forest map by Cox regression analysis concerning integrated survival analysis of FDX1 expression in 33 cancer types **(A)** The expression of FDX1 is significantly related to OS in KIRC, LGG, HNSC, and CESC. **(B)** DFS, LIHC, and THCA show a significant correlation to FDX1. **(C)** KIRC, KIRP, and LGG are significantly related to FDX1 expression. **(D)** PFS of ACC, KIRC, LGG, MESO, and THCA is significantly related to FDX1 expression. (P<0.05 indicates that the expression of FDX1 is significantly correlated with the prognosis and survival of the patients. HR>1, with the increase of FDX1 expression, the risk of poor prognosis is higher).

**Figure4.**
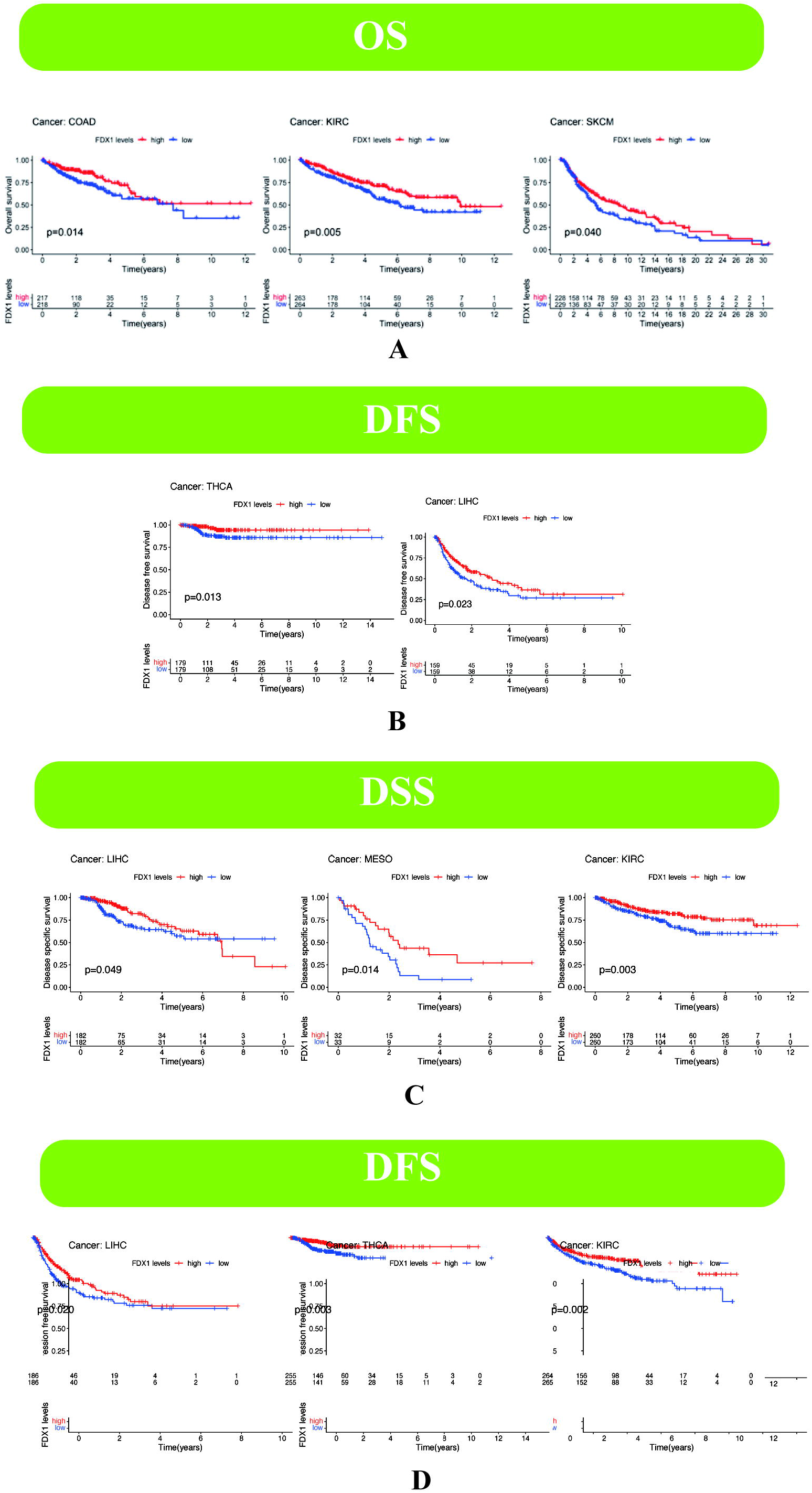
Low FDX1 expression is associated with poor survival in 6 tumors. **(A)**Correlation between FDX1 expression and OS in 3 cancers, including COAD(p=0.014), KIRC(p=0.005), SKCM(p=0.040) **(B)** Correlation between FDX1 expression and DFS in 3 cancers, including THCA(p= 0.013), LIHC(p= 0.023) **(C)** Correlation between FDX1 expression and DSS in 5 cancers patients, including LIHC(p= 0.049), MESO(p= 0.014), KIRC(p= 0.003) **(D)** Correlation between FDX1 expression and PFS in 5 cancers, including LIHC(p= 0.020), THCA(p= 0.003), KIRC(p= 0.002). p< 0.05 represents a significant difference.

Subsequently, GO term and KEGG pathway analyses after the normalization and preparation of transcriptome data from the TCGA Database were conducted to further understand the biological function of FDX1 in 11 cancers in which FDX1 expression correlates with patient prognosis. The curve peak upward indicated that GSEA results were positively correlated with FDX1 high expression. Finally, what we found is, that the biological processes, molecular functions, and cellular components closely related to FDX1 expression include ncRNA3 End, snRNA metabolic, positive regulation of t cell proliferation, integrator complex, odorant binding, keratin filament (Figure 5A). KEGG pathway analysis indicated five pathways significantly correlated with FDX1 expression were pathways in cancers, porphyrin and chlorophyll metabolism, cytosolic DNA sensing pathway, neuroactive ligand-receptor interaction, olfactory transduction, focal adhesion, regulation of actin cytoskeleton, chemokine signaling pathway, cytokine receptor interaction, nod like receptor signaling pathway, cell adhesion molecules cams, ECM receptor interaction, calcium signaling pathway (Figure 5B). Therefore, we speculated that FDX1 expression may be involved in the occurrence and development of various cancers through the oxidative metabolism pathway regulating the integrator complex.

**Figure5.**
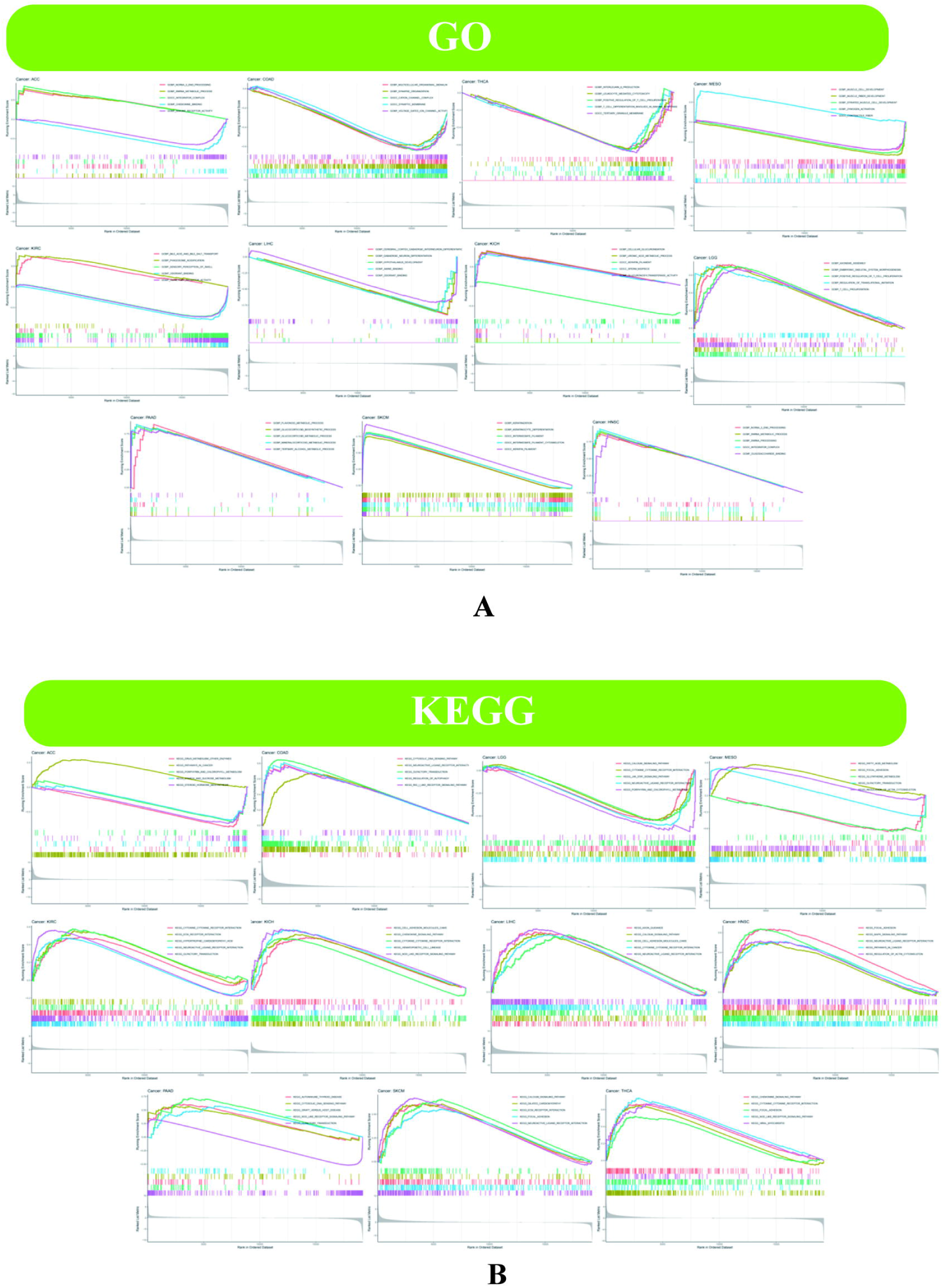
Enrichment analysis of FDX1 expression in 11 tumors, including ACC, COAD, HNSC, KIRC, LGG, PAAD, SKCM, KICH, LIHC, THCA, MESO. **(A)**GO analysis was performed in 11 tumors. **(B)**KEGG pathway analysis with FDX1 expression in 11 tumors. (The curve peak upward means the enriched pathway is positively correlated with FDX1, while the curve peaks downward negatively).

### Correlations of FDX1 expression levels with TMB and MSI

TMB and MSI are intrinsically related to the sensitivity of immune checkpoint inhibitors (ICI). The results of this study show that the expression level of FDX1 is related to the TMB of 11 types of cancer. Moreover, the expression of FDX1 was positively correlated with TMB in LGG, ESCA, STAD, UCEC, and PRAD, but negatively correlated with TMB in KIRC, LUAD, THCA, KICH, and THYM (Figure 6A). In another 9 types of tumors, FDX1 expression was related to MSI. Besides, the expression of FDX1 was positively correlated with MSI in DLBC, STAD, KIRC, UCEC, and HNSC, while negatively correlated with MSI in LUAD, LUSC, PAAD, and ACC (Figure 6B).

**Figure6.**
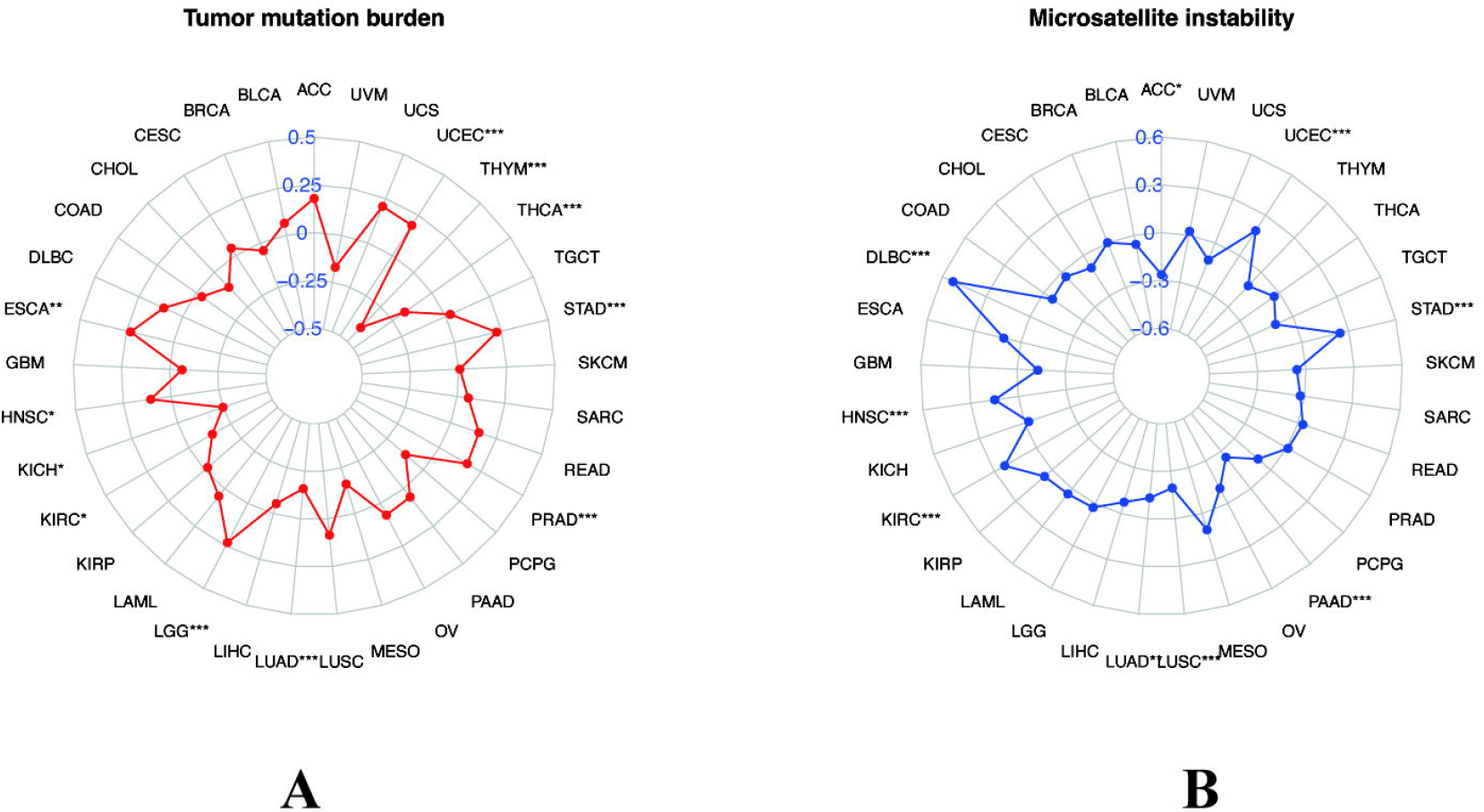
Radar map of the correlations of FDX1 expression levels with TMB and MSI. **(A)** Correlations of FDX1 Expression Levels with TMB **(B)** Correlations of FDX1 Expression Levels with MSI. Spearman Correlation test, p<0.05 was considered significant, *p < 0.05, **p < 0.0l, ***p < 0.001

### FDX1 expression with TME and immune cell infiltration analysis in pan-cancers

TME consists of tumor cells and non-tumor components which are mainly composed of stromal and immune cells. Therefore, the higher the stromal and immune cell scores in tumor tissue, the lower the purity of the corresponding tumor. The figure accounts for which there exists a positive correlation between FDX1 level and immune cell scores in PCPG, SARC, and LGG (Figure7 D-F), stromal scores in TGCT and LGG (Figure7 G-H), while negative and stromal and immune cell scores in LGG and stromal scores in STAD (Figure7 A-C). Besides, the analysis results of immune cell infiltration indicate that FDX1 expression is correlated with the immune cell infiltration of the 3 tumors, including ACC, KICH, and UVM. Among them, the infiltration of eosinophils and monocyte is related to the high expression of FDX1 in ACC and KICH, while, in ACC, KICH, and UVM, the infiltration of T cells CD8, Macrophages M0, and Monocytes the low expression of FDX1(Figure8). In addition, the results of co-expression FDX1 with immune cell genes illustrate that the immunosuppressive gene TGEBR1 in UVM, immunoregulatory gene CD86 in LGG, and MHC gene HLA-DQA2 in KICH are the strongest positive correlation gene with FDX1(Figure9). Above all the research confirm that gene levels at immune checkpoints are significantly elevated in cancers that were positively correlated with FDX1 expression, which prompted us to wonder whether FDX1 expression is related to immunotherapy in cancer patients.

**Figure 7.**
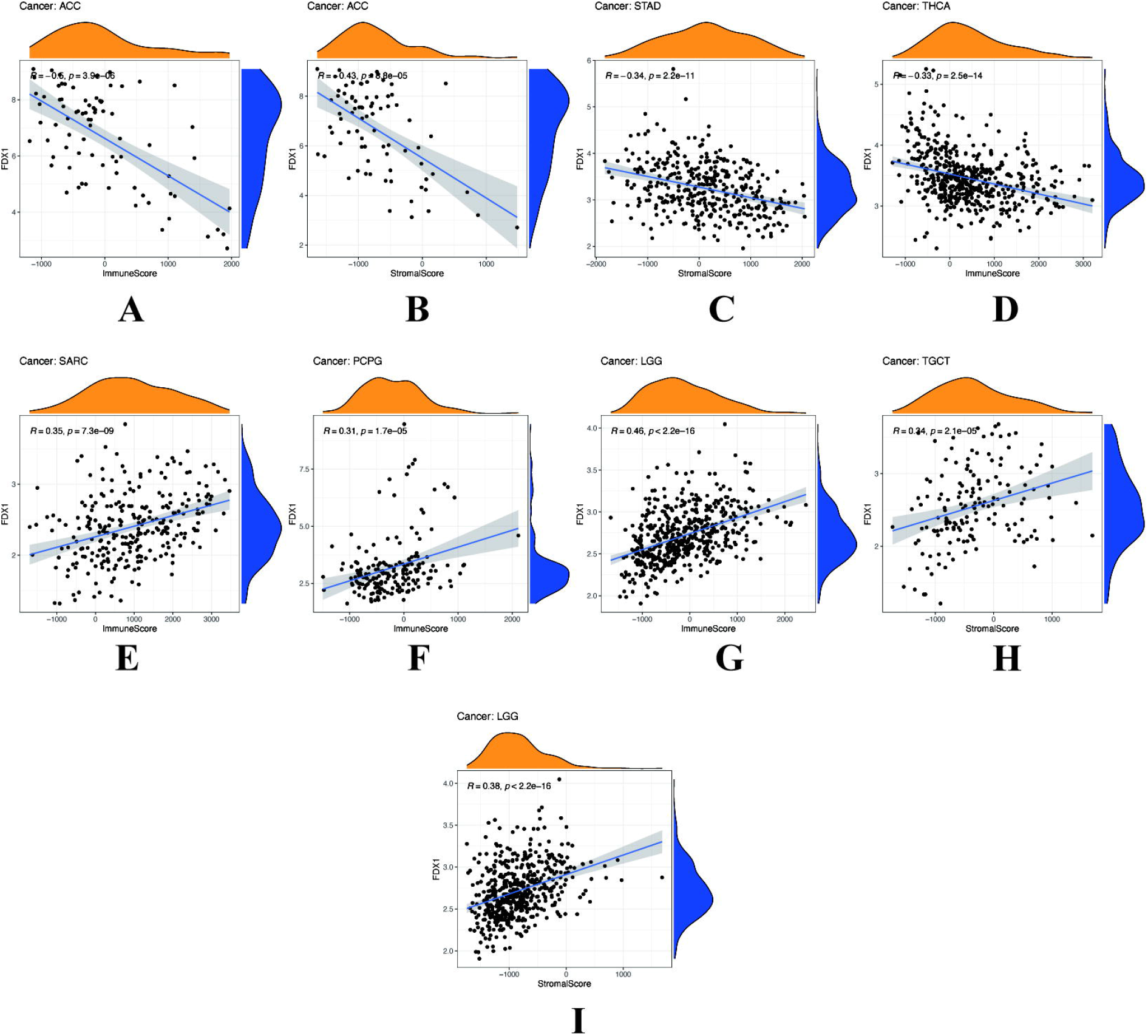
Correlation between FDX1 expression and the tumor microenvironment in 7 cancers, including ACC, STAD, THCA, SARC, PCPG, LGG, and TGCT. (P<0.05 indicates that the expression of FDX1 is significantly correlated with t the tumor microenvironment. R>0, with the increase of FDX1 expression, the estimated score is higher).

**Figure 8.**
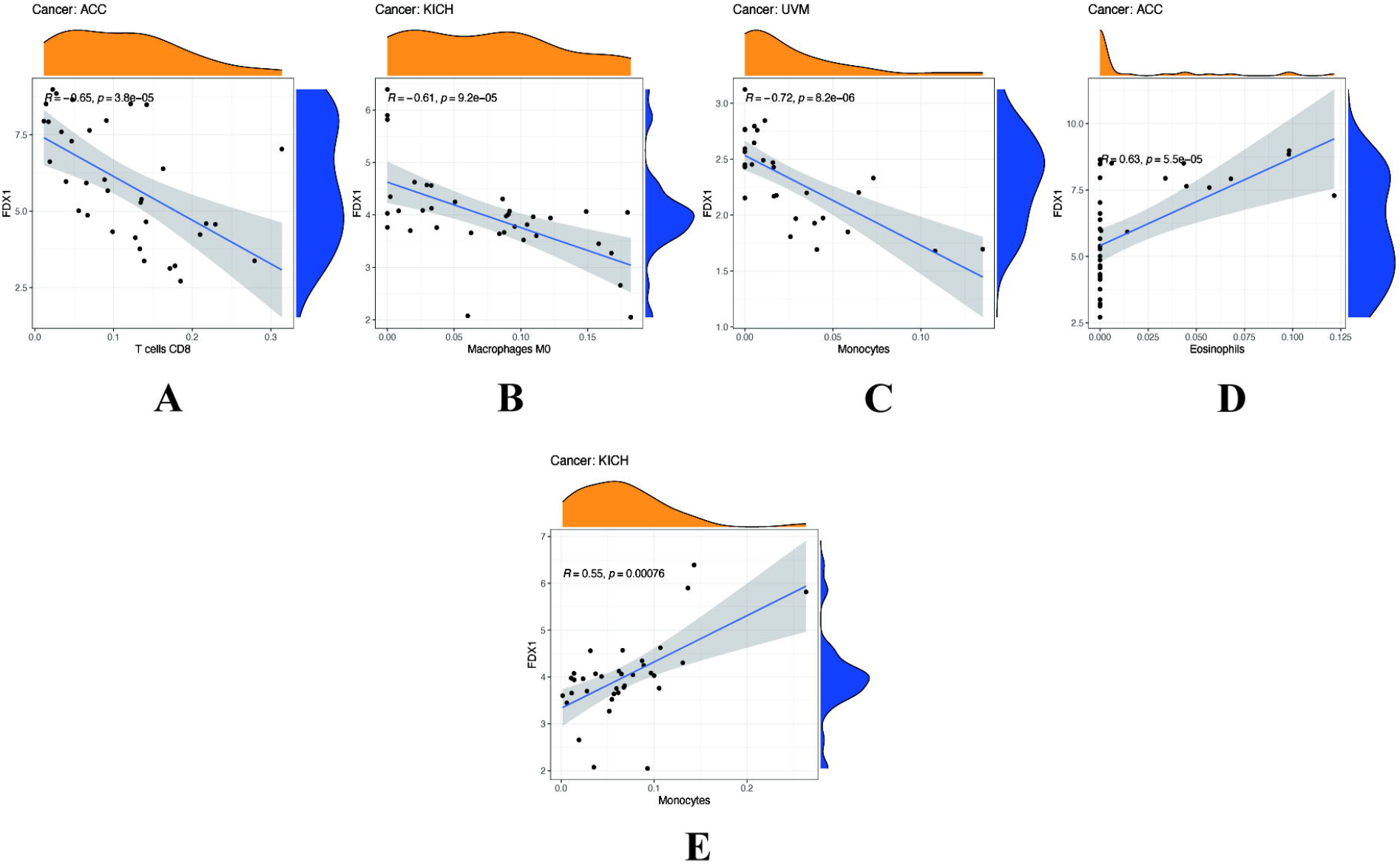
Correlation between FDEX1 expression and immune infiltration in ACC, KICH, UVM. (P<0.05 indicates that the expression of FDX1 is significantly correlated with immune infiltration. R>0, with the increase of FDX1 expression, the level of immune cells is higher).

**Figure9.**
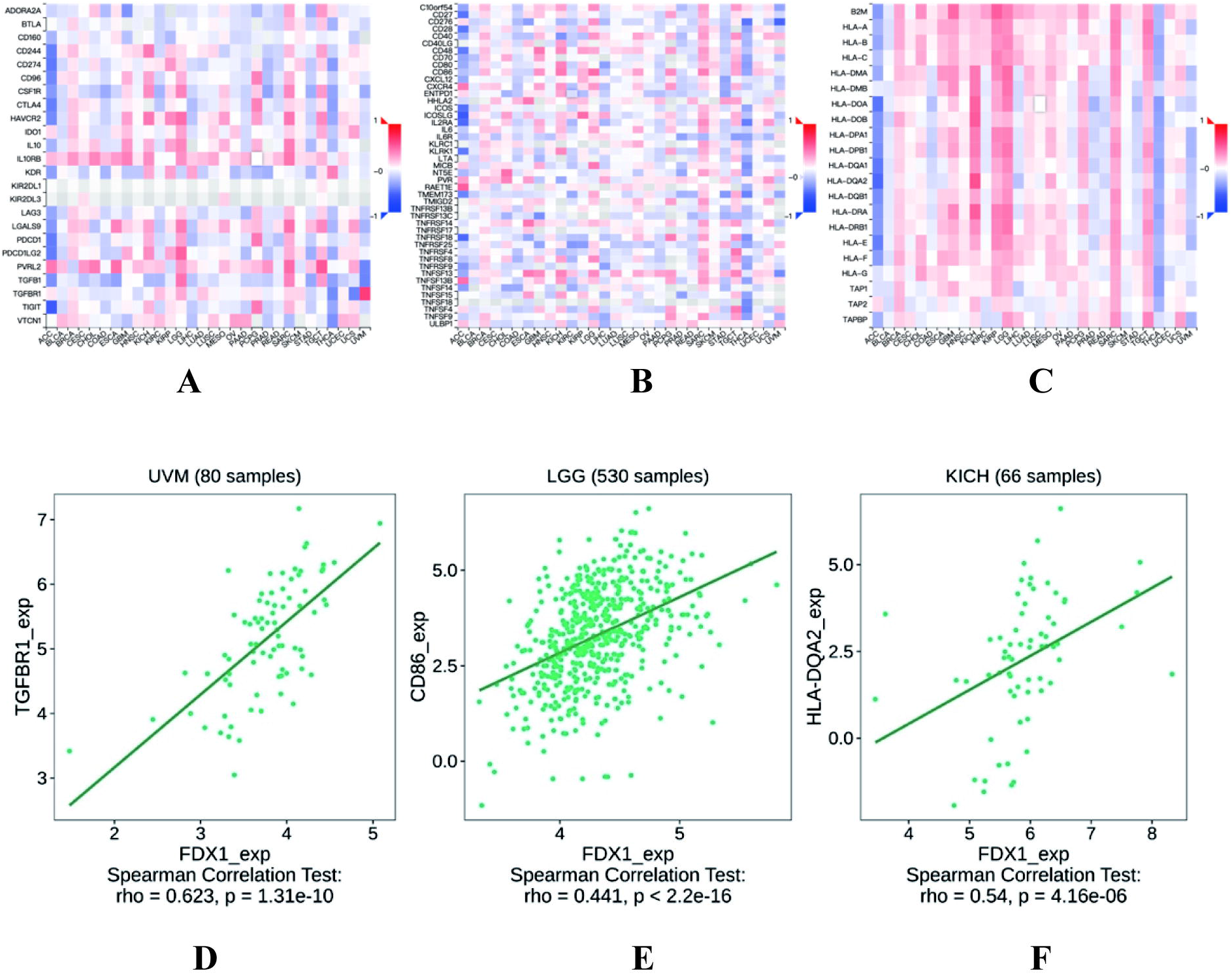
Correlation between FDX1 and immune gene expression in 33 cancers. **(A)** The heat map of gene co-expression of FDX1 and immunoinhibitory **(B)** The heat map of gene co-expression of FDX1 and immunostimulatory **(C)** The heat map of gene co-expression of FDX1 and MHC molecule **(D)**Correlation between immunosuppressive gene TGEBR1 and FDX1 expression in UVM **(E)**Correlation between immunoregulatory gene CD86 and FDX1 expression in LGG **(F)**Correlation between MHC gene HLA-DQA2 and FDX1 expression in KICH. (Spearman Correlation test, p<0.05 was considered significant, rho>0, with the increase of FDX1 expression, the immune gene expression is higher).

## Discussion

FDX1 expression can promote the death of tumor cells ^[10]^. What’s more, the pathway that FDX1 induces tumor cell death has been illuminated in the recent research titled “Copper ionophore–induced cell death” published in the journal Science^[17]^. Copper death has been proved to be closely related to the progression of many tumors^[18-21]^, in which the accumulation of copper in cells results in an overall reduction in proteins involved in mitochondrial respiration, reduced protein lipoylation, and reduced levels of Fe-S cluster proteins. According to the evidence mentioned, there exists a hypothesis that in these cancers FDX1 serves as a protective gene FDX1 launch the mechanism of copper death in the tumor cell. Therefore, it is essential to study the role of FDX1 in various cancer types and testify to the guess by related tumor cell experiments in prospective research.

Conversely, it is a pity that there is still a lack of extensive analysis of FDX1. Given this situation, in our study, we comprehensively analyzed the expression of FDX1 in multiple tumors based on various data sets and explored the correlation between FDX1 expression and the age, gender, tumor stage, prognosis, and tumor microenvironment. Furthermore, the biological functions and related pathways involving FDX1expression in the development of tumors also were discussed in depth. Finally, we demonstrated the value of FDX1 as a prognostic biomarker in a variety of tumors and its role in immunotherapy.

First of all, we conducted the expression of FDX1 in tumor and normal samples across disparate cancers. Results testify that FDX1 was expressed in tumor samples of all cancers. However, FDX1 levels vary among types of cancer, which may be due to the different levels of copper and lipoylated components in a variety of tumor tissues, as well as the different dependence of tumor cells on mitochondrial respiration^[22]^. Besides, there was a significant difference in FDX1 expression between tumor and normal samples in 17 cancers, among which FDX1 was highly expressed, including UCEC, STAD, and lowly expressed, including BRCA, CHOL, COAD, GBM, HNSC, KICH, KIRC, KIRP, LIHC, LUAD, LUSC, PCPG, READ, SARC, SARC THCA. The above results are consistent with previous studies which proved that the expression levels of FDX1 in the tumor tissues of ACC, KICH, KIRC, KIRP^[11]^, LIHC^[12]^, and LUAD^[10]^ were lower than those in the normal tissues.

Secondly, copper-induced cell death with FDX1 as a key regulator has been shown to inhibit tumor progression^[19]^, which led us to speculate that FDX1 expression is of great significance in the clinical progression of tumors. What’s striking is that the results of which FDX1 level decreased gradually in KICH, LIHC, and THCA as the tumor progressed from stage I to stage III verified our hypothesis, indicating that it may be possible to regulate FDX1 expression as a target to induce tumor cell apoptosis. Meanwhile, the higher survival rate of 11 types of cancer patients with high expression of FDX1 further confirmed the protective effect of the high level of FDX1 on tumors, which illustrated the value of FDX1 as a prognostic biomarker for various cancers. The GSEO results, which involved biological functions and pathways related to metabolism in the low-expression FDX1 group, revealed the mechanism by which downregulation of FDX1 expression significantly affected metabolism. Previous experiments^[10]^ also have proved that FDX1 knockdown mainly promotes glycolysis and fatty acid oxidation, changes amino acid metabolism, and has a potential carcinogenic effect. These phenomena suggest that FDX1 works mainly by influencing the metabolic environment of tumor cells, which may enhance the immunotherapy effect of cancer. In addition, we found that FDX1 expression is different among cancer patients of different ages. For example, in ESCA, HNSC, UCEC, BRCA, AND KIRC, FDX1 expression level is higher in patients over 60 years old and lower in patients less than 60 years old, which can guide us to select different immunotherapy programs for different age groups to some extent.

Thirdly, TMB, as a promising biomarker for predicting generalized cancer^[23]^, plays an important role in guiding immunotherapy for cancer patients^[24]^. For example, studies have reported that TMB as a biomarker can significantly improve the cure rate of non-small cell lung cancer^[25]^ and colorectal cancer^[26]^. In addition, high-frequency MSI is an independent predictor of clinical characteristics and prognosis in patients with colorectal cancer, as well as an important biomarker^[27]^. In this study, FDX1 expression was correlated with TMB in 11 cancers and MSI in 9 cancers, suggesting that FDX1 expression may affect the efficacy of ICI therapy by affecting TMB and MSI, which may provide some new references for the prognosis of immunotherapy in the future. Based on the studies that have been done so far and our findings, we speculated that among the cancers with a positive correlation between TMB and FDX1 expression, those with high expression of TMB, MSI, and FDX1 had a better prognosis after Cancer immunotherapy (CIT).

Fourthly, on the one hand, studies ^[28-34]^ have given credible proof that the specific metabolism in cancer cells produced a hypoxic, acidic, nutrient-depleted TME that is not conducive to anti-tumor immune response, promoting tumor growth and preventing the generation of the effective anti-tumor immune response. Besides, since fatty acid metabolism has a significant influence on TME and immunotherapy^[6, 35]^, while the low expression of FDX1 mainly affects substance metabolism (the results of the GSEA), we further explored the relationship between FDX1 expression and tumor microenvironment and immune cell infiltration. Finally, we found that FDX1 expression was positively correlated with stromal and immune cell content in PCPG, SARC, LGG, and TGCT, which illustrates that patients in these cancers are more responsive to immunotherapy. In addition, immune cells can antagonize or inhibit the occurrence and development of tumors, so tumor-infiltrating immune cells play an important role in immunotherapy^[36]^. For instance, positive correlations between FDX1 and eosinophils and monocyte in ACC and KICH manifested that high expression of FDX1 affected immunotherapy by promoting their invasion in tumor tissues, which mainly attributed to the formidable ability that eosinophils affect local immunity and tissue remolding during homeostasis and disease^[37]^. Currently, there is a lot of evidence that eosinophils present direct or indirect antitumor activity in TME, which is closely related to the prognosis of cancer patients and can be used as a potential cell target for ICB therapy in the future^[38-40]^. In addition, the difference in the infiltration of monocytes with high expression of FDX1 between UVM and KICH may be due to the heterogeneity of monocytes, which homogeneous populations develop into heterogeneous cell systems, exhibiting different functions in response to diverse stimulus^[41]^. One characteristic of the successful immune response is the production of CD8 + memory T cells. What’s more, Memory T cell formation is completely dependent on fatty acid oxidation and mitochondrial oxidative metabolism. In contrast, effector T cell response requires glycolysis and glutamine decomposition ^[42]^, which is consistent with our findings that CD8 + T cells were highly invasive in ACC with low FDX1 expression. Therefore, with the gradual accumulation of fatty acids in TME, effector T cells will be induced to senescence, promoting tumor progression. As for the finding of a negative correlation between Macrophages M0 and FDX1 expression, relevant studies have also reported at present that higher M0 macrophage counts are associated with poor prognosis in cancer patients^[43]^. All in all, we confirmed that high expression of FDX1 could promote the invasion of the relevant immune cells in the process of inhibiting tumor progression according to the relationship between fatty acid metabolism, immune cells, and FDX expression. This may provide vital ideas for finding new targets for immunotherapy.

On the other hand, a large number of studies have reported that the proliferation, metastasis, and recurrence of tumor cells are closely related to the immune escape of tumor^[44, 45]^. Therefore, the dual approach of eliminating immunosuppressive factors and looking for new biomarkers is necessary to achieve successful cancer treatment for the breakthrough of tumor immunotherapy. The co-expression of FDX1 with immune genes indicated that immunosuppressive gene TGEBR1 in UVM, immunoregulatory gene CD86 in LGG, and MHC gene HLA-Dqa2 in KICH were most correlated with the high expression of FDX1, which hints that we may be able to use these related immune genes as targets for immunotherapy. Furthermore, the downregulation of fatty acid oxidation by enhancing the level of FDX1 can reduce the number and function of Treg cells^[6]^, which can effectively fight tumor progression^[44]^. To sum up, FDX1 can be used as a new biomarker for tumor immunotherapy prognosis.

## Conclusion

In summary, our first extensive cancer analysis of FDX1 showed differential expression of FDX1 between tumors and normal tissues and revealed a correlation between FDX1 expression and clinical prognosis and immune cell infiltration in various types of cancer, suggesting that FDX1 may be a potential biomarker of immunotherapy response. Future studies of FDX1 expression and immune cell infiltration in different cancer populations can be further prospectively and experimentally investigated. Furthermore, FDX1 can be used as an independent prognostic factor for many tumors. For different tumors, the level of FDX1 expression will bring diverse prognostic results, so the specific role of FDX1 in each cancer needs to be further studied. In addition, FDX1 expression is related to TMB, MSI, and TME. Its effect on tumor immunity also varies by tumor type. These findings may help clarify the role of FDX1 in tumor genesis and development, which provide a reference for the realization of more accurate and personalized immunotherapy in the future.

## Data availability statement

The original contributions presented in the research are included in the article/Supplementary Material, more inquiries can be directed to the corresponding author.

## Ethics statement

As this work benefited from the database of TCGA, informed consent was not applicable.

## Author contributions

Qingqing Zhang wrote the article. Ping Wang, Nanyang Liu, Zhengyu Xu, and Yichen Wang participated in the preparation of the manuscript, construction of tables and citations of references, and the critical revision of the literature. All authors contributed to the article and approved the submitted version.

## Abbreviations

ACC: adrenocortical carcinoma
AR: androgen receptor
BLCA: bladder urothelial carcinoma
BRCA: breast invasive carcinoma
CESC: cervical squamous cell carcinoma and endocervical adenocarcinoma
CHOL: cholangiocarcinoma
COAD: Colon adenocarcinoma
CRC: colorectal cancer
DFI: disease-free interval
DLBC: lymphoid neoplasm diffuse large B-cell lymphoma
DSS: disease-specific survival
ESCA: esophageal carcinoma
ESTIMATE: Estimation of stromal and Immune cells in Malignant Tumors using Expression data
HNSC: head and neck squamous cell carcinoma
KICH: kidney chromophobe
KIRC: kidney renal clear cell carcinoma
KIRP: kidney renal papillary cell carcinoma
LGG: brain lower-grade glioma
LIHC: liver hepatocellular carcinoma
LUAD: lung adenocarcinoma
LUSC: lung squamous cell carcinoma
MESO: mesothelioma
MSI: microsatellite instability
OS: overall survival
PAAD: pancreatic adenocarcinoma
PCPG: pheochromocytoma and paraganglioma
PFI: progression-free interval
PRAD: prostate adenocarcinoma
READ: rectum adenocarcinoma
SARC: sarcoma
SKCM: skin cutaneous melanoma
STAD: stomach adenocarcinoma
TCGA: The Cancer Genome Atlas
TGCT: testicular germ cell tumors
THCA: thyroid carcinoma
THYM: thymoma
TMB: tumor mutational burden
TME: tumor microenvironment
UCEC: uterine corpus endometrial carcinoma
UCS: uterine carcinosarcoma
UVM: uveal melanoma

